# Spatial variation in water loss predicts terrestrial salamander distribution and population dynamics

**DOI:** 10.1101/004986

**Authors:** W.E. Peterman, R.D. Semlitsch

## Abstract

Many patterns observed in ecology, such as species richness, life history variation, habitat use, and distribution have physiological underpinnings. For many ectothermic organisms temperature relations shape these patterns, but for terrestrial amphibians, water balance may supersede temperature as the most critical physiologically-limiting factor. Many amphibian species have little resistance to water loss, which restricts them to moist microhabitats and may significantly affect foraging, dispersal, and courtship. Using plaster models as surrogates for terrestrial plethodontid salamanders, we measured water loss under ecologically-relevant field conditions to estimate the duration of surface activity time across the landscape. Surface activity time was significantly affected by topography, solar exposure, canopy cover, maximum air temperature, and time since rain. Spatially, surface activity times were highest in ravine habitats and lowest on ridges. Surface activity time was a significant predictor of salamander abundance, as well as a predictor of successful recruitment; the probability of a juvenile salamander occupying an area with high surface activity time was two times greater than an area with limited predicted surface activity. Our results suggest that survival, recruitment, or both are demographic processes that are affected by water loss and the ability of salamanders to be surface active. Results from our study extend our understanding of plethodontid salamander ecology, emphasize the limitations imposed by their unique physiology, and highlight the importance of water loss to spatial population dynamics. These findings are timely to understanding the effects that fluctuating temperature and moisture conditions predicted for future climates will have on plethodontid salamanders.

## Introduction

An organism’s physiology is dynamically related to its environment; physiology dictates the habitats that are occupied and behaviour within these habitats, while the environment can affect physiological performance and subsequently, ecological performance (Huey 1991). In concert with the environment, physiology can affect an organism’s performance at the local scale (Brewster et al. 2013), life history at a regional scale (Kearney 2012), and dictate limits on distribution (Buckley et al. 2010; Gifford and Kozak 2012; Kearney and Porter 2009). Further, potential responses to habitat or climate change can be modelled mechanistically by incorporating physiology (Kearney et al. 2008; Keith et al. 2008; Sinervo et al. 2010). The role of physiology is especially evident in ectothermic organisms, with the preponderance of emphasis being placed on thermal aspects of behaviour, physiology, and life history evolution (Angilletta 2009; Angilletta et al. 2004).

Although not independent of temperature and metabolic processes, water balance is another critical physiological characteristic that weighs heavily on the behaviour, distribution, and ecology of terrestrial taxa, especially amphibians (Tracy et al. 2010; Wells 2007). The skin of most amphibians provides little to no resistance to water loss (Spight 1968; Spotila and Berman 1976), even when the atmosphere is near saturation (Adolph 1932). All terrestrial amphibians must manage their hydric relations, but it is particularly critical for woodland salamanders of the genus *Plethodon.* These salamanders are unique among terrestrial vertebrates in that they are lungless and respire cutaneously by diffusion (Whitford and Hutchison 1967). As a consequence, plethodontid skin must remain moist and permeable to facilitate gas exchange, but these requirements impose physiological and ecological limitations. Because of its permeability, the skin of plethodontid salamanders loses water at a rate that is nearly identical to a free water surface of equivalent surface area (Peterman et al. 2013; Spotila and Berman 1976). Uninhibited water loss impinges upon salamander activity, potentially limiting foraging, dispersal, and reproductive efforts. Terrestrial plethodontid salamanders spend the majority of their life under ground or sheltered by cover objects such as rocks and logs on the ground surface (Petranka 1998). Surface activity and foraging of salamanders is greatest under moist conditions (Grover 1998; Keen 1979; Keen 1984), and the duration of time spent foraging is directly tied to water balance (Feder and Londos 1984). To minimize water loss, salamanders are predominantly nocturnal, and are generally associated with cool, moist microhabitats (Heatwole 1962; Peterman and Semlitsch 2013; Spotila 1972).

From a physiological perspective, four measurements are needed to predict the duration of salamander surface activity: salamander mass (used to calculate surface area; Whitford and Hutchison 1967), air temperature, relative humidity, and wind speed (Feder 1983). These factors can be used to predict that surface activity will be greatest for large salamanders when humidity is high, temperatures are cool, and there is no wind. Ecologically, this means that adults may have an advantage over juveniles in being able to sustain prolonged surface activity due to their lower surface area to volume ratio, and microclimate variation produced by landscape features such as topography may profoundly affect surface activity times by modulating temperature, wind, and humidity. Limited surface activity may limit foraging time, and consequently affect individual growth and reproduction. Dispersal may also be curtailed, reducing gene flow among local populations.

Despite the intuitive effects that hydric constraints impose on terrestrial plethodontid salamander activity time, habitat use, and population dynamics, direct tests of these processes have been limited. Within a controlled laboratory setting, Feder and Londos (1984) found that a stream salamander (*Desmognathus ochrophaeus*, Cope) would abandon foraging in dry air twice as quickly as in moist air (3.8% vs. 7.5% loss of body mass, respectively). Grover (1998) experimentally demonstrated that surface activity of salamanders, especially juveniles, increased with increased soil moisture. Peterman and Semlitsch (2013) found that terrestrial salamander abundance was greatest in dense-canopy ravines with low solar exposure and high moisture, and found evidence of differential reproductive success related to these landscape features. Effects on population dynamics have indirectly been observed through variation in egg production. Grover and Wilbur (2002) found that salamanders in high moisture conditions produced more eggs, and both Milanovich et al. (2006) and Maiorana (1977) found annual fecundity to increase with precipitation. These findings suggest that wetter conditions may accommodate increased surface activity and foraging, increasing the energy available to be allocated to reproduction.

By incorporating physiology with spatial and temporal climate variation, mechanistic biophysical models are capable of accurately predicting the distribution (Kearney and Porter 2009), biotic interactions (Buckley and Roughgarden 2005; Gifford and Kozak 2012), and life history variation (Kearney 2012; Tracy et al. 2010) of species. To encompass spatial heterogeneity, most of these studies cover broad geographical or elevational ranges. However, environmental gradients can occur over significantly smaller spatial scales in topographically complex landscapes (Bennie et al. 2008; Chen et al. 1999). Further, fine-scale variation in microclimate can affect species occurrence, population dynamics, and resilience to changing climatic conditions, especially in species with low vagility (Antvogel and Bonn 2001; Peterman and Semlitsch 2013; Scherrer and Körner 2011; Weiss et al. 1988). Although the importance of fine-scale microclimatic variation is well-understood (Huey 1991), most analyses of physiological processes have not been spatially explicit.

The objective of our study was to explicitly test, for the first time, the hypothesis that water balance is a limiting factor for terrestrial salamanders (Feder 1983). Specifically, that spatial variation in water loss and surface activity time affects the distribution of salamanders as well as population dynamics across the landscape. We model physiological landscapes describing fine-scale spatial variation in water loss rates for a terrestrial plethodontid salamander, *Plethodon albagula* (western slimy salamander), and then convert these rates to potential surface activity times. In calculating rates of water loss surface activity time we seek to (1) determine the landscape and environmental factors influencing spatial variation in water loss in a topographically complex landscape, (2) determine whether salamander distribution on the landscape can be predicted by the physiological limitations imposed by water loss and activity time, and (3) assess the effects of surface activity time on spatial population dynamics. We hypothesized that rates of water loss would be dependent upon both topographical landscape features as well as climatic conditions. Specifically, we predicted that topographic complexity would result in heterogeneous water loss rates across the landscape and that ravine habitat with low solar exposure would have the lowest rates of water loss. Temporally, we predicted that abiotic factors such as time since rain, air temperature, and relative humidity would significantly affect daily and seasonal patterns of water loss. Because Peterman and Semlitsch (2013) found salamander abundance to be greatest in sheltered ravine habitats and lowest on ridges, we hypothesized that spatial patterns of water loss would corroborate these patterns with ravines exhibiting low rates of water loss and ridges high rates of water loss. We also hypothesized that water loss, as an integrated measure of the landscape and climate, would significantly predict the spatial distribution of salamander abundance. Lastly, as a mechanism limiting population growth, we hypothesized that evidence of successful reproduction would be greatest in ravines with lower rates of water loss.

## Materials and methods

### STUDY SPECIES

*Plethodon albagula* (western slimy salamander, Grobman) are a large plethodontid salamander of the *P. glutinosus* species complex that live in forested habitats throughout the Ozark and Ouchita mountains of Missouri, Arkansas, eastern Oklahoma, and northeastern Texas, USA (Highton 1989). Within these forested habitats, salamander abundance is greatest in moist, forested ravines (Peterman and Semlitsch 2013). Surface activity varies seasonally, with peak activity occurring in spring and to a lesser extent during autumn (Milanovich et al. 2011); terrestrial plethodontid salamanders generally seek subterranean refuge during hot, dry summer conditions (Taub 1961).

### PLASTER MODELS

We assayed water loss across the landscape using cylindrical plaster of Paris models (hereafter “replicas”) as analogues for live salamanders. Plaster replicas were made following methods described by Peterman et al. (2013), and had surface areas equivalent to adult- and juvenile-sized salamanders that were 7.25 g and 2.25 g, respectively. Previous research has shown these replicas lose water linearly and at rates equivalent to similarly-sized salamanders (Peterman et al. 2013). Models were cured in a drying oven for 24 h at 70°C, and then weighed to the nearest 0.01 g on a portable digital balance (Durascale, My Weigh, Vancouver, BC). Prior to deployment, all replicas were soaked in water for at least four hours; replicas were deployed within one hour of sunset, and retrieved within one hour of sunrise.

Replicas were deployed at Daniel Boone Conservation Area (DBCA; Fig. 1) along 250-m long transects, spaced at approximately 50-m intervals (n=18 transects; 108 locations). Locations of replica deployment were marked in the field using a handheld GPS (Garmin 62sc, Olathe, Kansas, USA) with multiple locations being taken until the estimated precision was ≤3 m. Replicas were deployed in both spring (8 April–8 May 2012) and summer (15 August–28 August 2012). At each location, adult- and juvenile-sized replicas were deployed under the leaf litter, and another pair was deployed on top of the leaf litter. Because the focus of this study was the effects of landscape and climate features on water loss, all replicas were housed within cylindrical cages made of 3 mm hardware cloth to prevent replicas from coming in direct contact with leaf litter or soil, which could have confounding effects on water loss rates (Peterman et al. 2013). Each replica was weighed with the portable digital balance upon deployment and retrieval.

**Figure 1.**
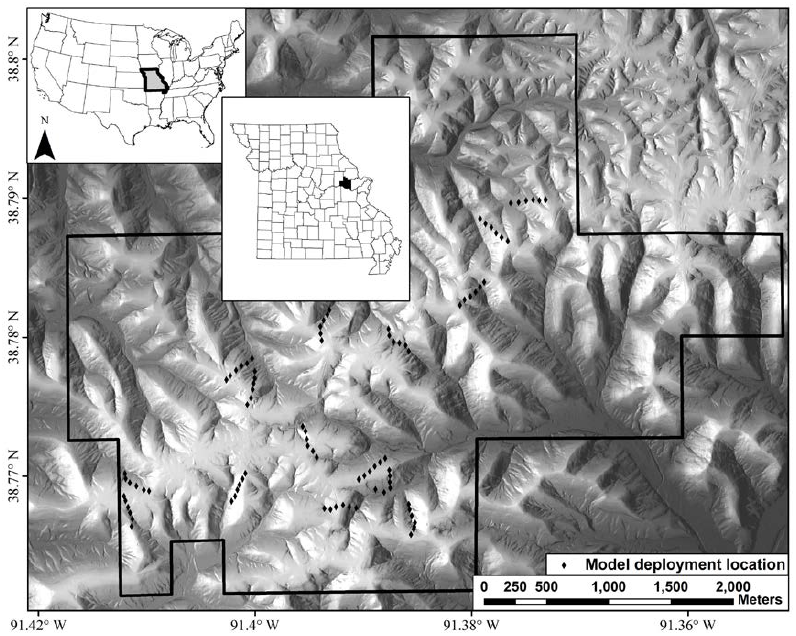
Locations of plaster replica deployment at Daniel Boone Conservation Area, Missouri, USA. Replicas were deployed at six locations separated by 50 m along each transect, and there were 18 transects (n = 108 locations).

**Figure 2.**
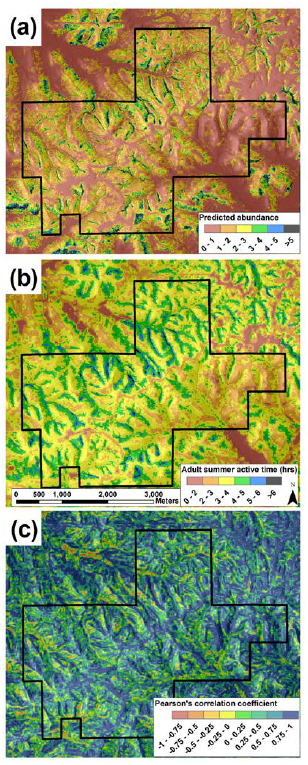
Maps of (a) predicted salamander abundance, (b) summer surface activity time (hrs) estimated from adult-sized plaster replicas, and (c) spatial Pearson’s *r* correlation values. There is generally very high, positive correlation between estimated abundance and surface activity time; as activity time increases, so does the predicted abundance of salamanders.

### SPATIAL AND TEMPORAL COVARIATES

Spatial covariates used in this analysis were calculated in ArcGIS 9.3 (ESRI, Redlands, CA, USA) and are described in detail in (Peterman and Semlitsch 2013). Previously, these covariates were used to predict the spatial distribution of abundance of *P. albagula* (see details below). In the current study we assessed the effects of topographic position (TPI), topographic wetness index (TWI), potential relative radiation (PRR), and distance from stream. These variables have a resolution of 3 m, and were derived from 1/9 arc second National Elevation Dataset (http://seamless.usgs.gov/products/9arc.php). Canopy cover was also estimated at DBCA using the normalized difference vegetation index (NDVI), which was calculated from cloud-free Landsat 7 satellite images of our study area taken on 15 June, 20 July, 9 August 2012 (http://glovis.usgs.gov/). A mean NDVI was calculated by averaging these days together. The resolution of the NDVI layer was 30 m, so it was resampled to a resolution of 3 m. Because the majority of our spring trials were conducted prior to full leaf-out, NDVI was not included in the spring models. For this analysis, we used time since rain, maximum overnight humidity, and maximum temperature of the previous day as temporal climatological covariates. These data were collected from the Big Spring weather station (<u>http://www.wunderground.com</u>), which is located 8 km west of DBCA. For extrapolating our model to the entire DBCA landscape, we determined averages for these measures in spring (1 April–31 May) and summer (1 June–31 August) from data collected 2005–2012.

### STATISTICAL ANALYSES

For each replica we calculated the proportion of water lost per hour (*proportion loss = [deployed mass - retrieved mass] / [deployed mass - dry mass] / time deployed*), which became our dependent variable. For this analysis, we did not have competing *a priori* hypotheses concerning the factors that would affect water loss, but rather, we were interested in fitting the best model possible to explain the spatial and temporal patterns of water loss in our plaster replicas. As such, we did not conduct extensive model selection on parameters to include or exclude from each model, but instead fit a small number of meaningful parameters to each model. Our modeling work flow proceeded as follows. We first divided our data by replica size and season (size-season) to create four independent data sets (juvenile-spring, juvenile-summer, adult-spring, and adult-summer). We then assessed the correlation of each of our independent variables with each other, as well as their correlation with the dependent variable. If two variables had a Pearson’s correlation *r*≥0.70, we excluded the variable that had the lowest correlation with the dependent variable. Lastly, to limit complexity we did not include interactions of independent variables, and excluded variables that had *r*<0.10 correlation with the dependent variable. To account for heterogeneous variance in our data, we fit different variance structures to our data using ‘nlme’ in R (Pinheiro et al. 2013; R Core Team 2013; Zuur et al. 2009). Model selection was based on AIC (Akaike 1974). Using the model with the best-fit variance structure, we then tested different random effects parameterizations to account for the nested nature of our data (i.e. models within location, locations within transect, transects within date). The percent variance explained by our top model for each size-season combination was assessed using the marginal *R*^2^ measure of Nakagawa and Schielzeth (2013) and calculated with ‘MuMIn’ (Barton 2013). The marginal *R*^2^ describes the percent variation explained in the fixed effects model alone. The full list of variance structures and random effects parameterizations tested in model selection can be found in Appendix S1.

The fixed effects parameter estimates for the top size-season models were then used to predict water loss rates across the DBCA landscape. Replica position (under leaves or on the surface) was a factor in each model, so for each size-season combination, we calculated a surface and a leaf water loss estimate. For the remainder of this paper we consider salamander surface activity to be evenly divided between these two states (i.e. 50% surface, 50% under leaves). Therefore, to calculate a single size-season water loss rate, we averaged the model predictions from surface and leaf models. Because the main objective in this study is to demonstrate water loss as a limiting factor for terrestrial salamanders, we converted water loss rates to surface activity times (SAT). There is no empirical data describing the threshold of water loss when terrestrial plethodontid salamanders cease surface activity and seek refuge, and only one study has experimentally assessed this in a stream-associated salamander (Feder and Londos 1984). Previous studies have used 10% of body mass lost as the point at which salamanders stop foraging (Feder 1983; Gifford and Kozak 2012). For our study, we used 10% of total water lost as the threshold; SAT was calculated as the time (hrs) to 10% water loss. It should be noted that the proportion of a salamander’s body mass comprised of water decreases as mass increases (Peterman et al. 2013):

*Proportion Water = (−0.0168*live salamander wet mass (g)) + 0.8747*.

Ten percent mass loss for juvenile and adult salamanders of sizes equivalent to our replicas would result in 11.9% and 13.3% loss of water, respectively.

One of our objectives in this study is to determine how predicted SAT relates to the predicted spatial distribution of abundance. The methods and model used to predict salamander abundance across the landscape are described in detail by Peterman and Semlitsch (2013). Briefly, we surveyed 135 plots at DBCA that were spaced ≥75 m apart seven times in the spring of 2011. We fit binomial mixture models to our repeated count data using a Bayesian framework (Royle 2004). To account for imperfect observation of salamanders in space and time, we modeled salamander detection probability as a function of survey date, the number of days since a soaking rain event (rain ≥5mm), and temperature during each survey. After correcting for imperfect detection, abundance was modeled as a function of NDVI, TPI, TWI, and PRR. We then projected the fitted abundance model across the landscape to spatially represent the distribution of salamanders at DBCA.

We conducted Pearson product-moment correlation tests between the abundance estimates at the 135 survey plots from Peterman and Semlitsch (2013) and the spatial SAT predictions made in this study to get a point estimate correlation. We also assessed spatial patterns of correlation between SAT and abundance within ArcGIS using a moving window correlation (Dilts 2010) with a window size of 51 m (17 × 17 pixels). SAT is a physiological measure estimated from several of the same landscape covariates included in the abundance model of Peterman and Semlitsch (2013). To estimate the strength of SAT as a predictor of abundance, we re-ran the binomial mixture model of Peterman and Semlitsch (2013) in this study, but modeled abundance at each of the 135 survey plots solely as a function of SAT. Details of the model parameterization and settings can be found in Appendix S2.

Peterman and Semlitsch (2013) also used multistate models to identify a potential disconnect between reproductive effort (presence of gravid females) and realized recruitment (presence of juveniles). We generalize that analysis for this study to estimate the probability of juvenile and adult occurrence at each of the 135 plots surveyed by Peterman and Semlitsch (2013). We constructed multistate models using a conditional binomial parameterization in program PRESENCE v3.1 (MacKenzie et al. 2009). Models were fit separately for adult and juvenile salamanders, with three states being present in each model: (1) no salamanders present (site unoccupied); (2) salamanders present, but focal size class absent; (3) focal size class present, where the focal size class is either adult (snout-vent length [SVL] ≥55 mm; Milanovich et al. 2006) or juvenile (SVL <55 mm), respectively. As in the abundance model described above, we replaced the individual landscape covariates used by Peterman and Semlitsch (2013) with our integrated SAT measure. From this model we estimated the conditional probability of occurrence, which is the probability of a focal demographic group occurring at a site, given that a site is suitable to be occupied. Extended details of this analysis and model parameterization are in Peterman and Semlitsch (2013) and Appendix S2.

Lastly, we determined the mean SVL of salamanders observed at each of the 135 survey plots, and used a linear model to assess the relationship between SVL and SAT. Our objectives in re-analysing the data of Peterman and Semlitsch (2013) are to determine if SAT, as an integrated multivariate parameter, predicts abundance and occupancy of demographic groups, thereby providing a physiological mechanism for the effects of environmental gradients.

## Results

Correlations among independent variables revealed that TPI and distance from stream were highly correlated (*r* = 0.74), but TPI had a greater correlation with rate of water loss in the spring data sets, and distance to stream had a greater correlation in summer data sets. We also found TWI and maximum overnight humidity to have low correlation with water loss across all size-season combinations (*r*≤0.07), so these variables were not included in the mixed effects models. To account for heterogeneity within our data, an exponential variance structure was fit to both the juvenile and adult spring data, a combined identity-exponential variance structure was fit to the juvenile summer data, and an identity variance structure was fit to the adult summer data (Table 1). Random-effects fit to each model had both slopes and intercepts varying by covariates (Table 1). The average interval between rainfall events, as determined from the seven years of climate data, is 1.5 days (±1.98 SD) and 2.2 days (±2.85 SD) and the average daily maximum temperature is 22.5°C (±6.22) and 31.2°C (±4.00 SD) for spring and summer seasons, respectively. These seven-year mean estimates were used to make spatial predictions of water loss.

**Table 1.** Parameter estimates, standard errors (SE), and parameter significance from mixed effects model analyses of water loss rate for adult- and juvenile-sized plaster of Paris replicas.

| Fixed effects parameters | Spring water loss models |  |  |  |  |  | Summer water loss models |  |  |  |  |  |
| --- | --- | --- | --- | --- | --- | --- | --- | --- | --- | --- | --- | --- |
|  | Juvenile |  |  | Adult |  |  | Juvenile |  |  | Adult |  |  |
|  | Estimate | SE | P | Estimate | SE | P | Estimate | SE | P | Estimate | SE | P |
| Intercept | -3.968 | 0.931 | <0.0001 | -1.794 | 0.402 | <0.0001 | 16.075 | 6.874 | 0.022 | 6.360 | 4.903 | 0.201 |
| Position (surface) | 2.264 | 0.352 | <0.0001 | 1.155 | 0.184 | <0.0001 | 1.721 | 0.669 | 0.012 | 1.256 | 0.109 | <0.0001 |
| TPI<sup>a</sup> | 0.060 | 0.017 | 0.0006 | 0.004 | 0.007 | 0.546 | - | - | - | - | - | - |
| Dist from stream<sup>a</sup> | - | - | - | - | - | - | 0.007 | 0.002 | 0.005 | 0.007 | 0.002 | 0.006 |
| PRR | 0.000053 | 0.000060 | 0.378 | 0.000034 | 0.000027 | 0.206 | 0.000334 | 0.000110 | 0.003 | 0.000254 | 0.000107 | 0.022 |
| Max. Temp | 0.142 | 0.024 | 0.028 | 0.059 | 0.010 | 0.028 | -0.537 | 0.170 | 0.002 | 0.063 | 0.101 | 0.534 |
| Time since rain | 0.415 | 0.060 | 0.021 | 0.205 | 0.024 | 0.013 | 0.256 | 0.020 | <0.0001 | 0.167 | 0.013 | <0.0001 |
| NDVI<sup>b</sup> | - | - | - | - | - | - | 0.348 | 6.922 | 0.960 | -15.830 | 6.464 | 0.019 |
| $R^2_{GLMM(m)}$ | 89.79% | | | 93.68% | | | 98.69% | | | 82.90% | | |
| Linear model $R^2$ | 70.93% | | | 67.15% | | | 73.74% | | | 81.60% | | |
| Variance structure | Exponential (Max. Temp*position) |  |  | Exponential (Max. Temp*position) |  |  | Combined<br>Identity(position)<br>Exponential (Max. Temp*position) |  |  | Identity (position) |  |  |
| Random effects | ~1 + position\|date/transect/locale |  |  | ~1 + position\|date/transect/locale |  |  | ~1 + position\|date/transect/locale |  |  | ~1 + position\|locale |  |  |

<sup>a</sup> These parameters were correlated with each other; the parameter with the highest correlation with water loss rate was retained
<sup>b</sup> Spring models were deployed pre leaf-out, so canopy cover was not used as a predictor of water loss

Our final mixed effects models explained the majority of the variance in our data (*R*^2^_GLMM(*m*)_ = 82.90%–98.69%; Table 1). Notably, simple linear regression models that do not properly account for heterogeneity in variance or the nestedness of our sampling design described 67.15%–81.60% of variation in our data (Table 1). Plaster replica position was a significant predictor of water loss rate for both replica sizes in both seasons, with replicas on the surface losing 1.26%–2.64% more water per hour than adjacent replicas placed under leaves (Table 1). In the spring, water loss in juvenile replicas increased significantly with topographic position (TPI) meaning that water loss was greatest in ridge-like habitat and least in ravine-like habitats. In contrast, topographic position had no effect on adult replicas. Distance from stream had a significant effect on both juvenile and adult replica water loss in the summer with water loss rates increasing with distance from streams. Solar exposure (PRR) had no effect in the spring, but significantly increased rates of water loss in the summer (Table 1). The number of days since rainfall also significantly increased the rate of water loss in all size-season replicas. As anticipated, water loss increased with maximum temperature in the spring for both juvenile and adult replicas. Surprisingly, temperature had no effect on adult replica water loss in the summer, and had a negative effect on juvenile replica water loss (Table 1). Lastly, canopy cover, as measured by NDVI, was found to have no effect on juvenile replica water loss, but had a significant effect on adult replica water loss; as canopy cover increased, adult replica water loss decreased.

Spatially, there is extensive congruence among each size-season SAT map (Fig. S1), and correlations among these ranged from 0.62–0.95 (Table S1). The highest SAT are concentrated within ravine habitats, which are separated by ridges with lower SAT. Mean SAT on the landscape ranged from 1.94 hrs for juveniles in the summer, to 9.90 hrs for adults in the spring (Table 2). Paired t-tests revealed that juvenile SAT is significantly less than adult SAT in spring and summer, and that all SAT are significantly less in the summer (all tests *P*<0.0001). In general, the estimated SAT is 3 times longer in spring than summer, and is about 1.5 times longer for adults than juveniles, regardless of season (Table 2, Fig. S1). Correlations of predicted salamander abundance with size-season SAT at the 135 survey plots were also high (*r*=0.35–0.63; Table 2). Adult summer SAT had the highest correlation with predicted abundance (*r =*0.63), largely because of the significance of canopy cover in mitigating water loss (Table 1). Spatial similarities between predicted salamander abundance and adult summer SAT are evident (Fig. 3a–b); the correlation between abundance and SAT is generally highest in areas of low predicted abundance and low SAT (Fig. 3c).

**Figure 3.**
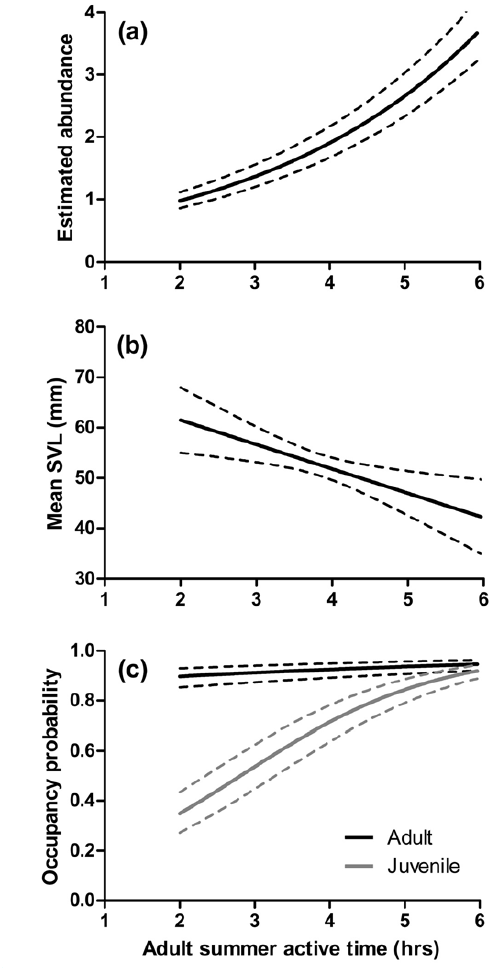
Relationship of estimated adult summer active time with (a) abundance; (b) mean SVL of salamanders; and (c) conditional probability of occurrence of adult and juvenile size classes. Dashed lines around the estimates represent 95% prediction intervals. Increased surface activity time resulted in more salamanders being present, and juvenile salamanders were more likely to be found in areas with higher surface activity times.

**Table 2.** Summary of estimated surface activity times (SAT*) and the correlation (Pearson’s *r*) of SAT with predicted abundance at Daniel Boone Conservation Area for juvenile- and adult-sized replicas in spring and summer seasons. Correlations are between predicted abundance from Peterman and Semlitsch (2013) and SAT from this study at 135 survey plots.

| Model | Surface active time (hrs) |  |  | Predicted abundance |
| --- | --- | --- | --- | --- |
| | Mean | SD | Range | Pearson's $r^{**}$ |
| Juvenile spring | 6.25 | 0.62 | 5.00–9.48 | 0.48 |
| Adult spring | 9.9 | 0.49 | 8.96–12.57 | 0.45 |
| Juvenile summer | 1.94 | 0.15 | 1.59–2.65 | 0.35 |
| Adult summer | 3.67 | 0.73 | 1.15–8.97 | 0.63 |

\* $SAT = 10 / \text{percent water lost}^{-hr}$
\*\* All correlations are significant at $P < 0.0001$

Because adult summer SAT had the highest correlation with abundance, we explored in more detail its relations with abundance, salamander size distribution, and probability of occurrence. We do note, however, that the other size-season models also had significant correlations with predicted abundance (*r*=0.35–0.48; Table 2). The binomial mixture model fit with predicted adult summer SAT as the sole independent variable in the abundance model fit the data well, and SAT had a significant effect on abundance, with abundance increasing as predicted SAT increased (Appendix S2; Fig. 3a). Further, we found that the mean SVL of salamanders observed at 88 of the 135 surveyed plots (n = 407 unique salamanders measured; Peterman and Semlitsch 2013) significantly increased as predicted SAT decreased (F_1, 86_ = 8.38; *P*=0.005; *R*^2^=0.089; Fig. 3b), suggesting that, on average, larger salamanders are found in areas with limited SAT. Similarly, we found that the conditional probability of juvenile salamander occupancy at the 135 surveyed plots, correcting for imperfect detection, significantly increased as predicted SAT increased (Appendix S2, Fig. 3c). In contrast, the conditional probability of adult occupancy was not significantly related to adult summer SAT (Fig. 3c), and there was little variation in predicted adult occupancy probability across the range of predicted SAT (adult conditional occupancy probability = 0.91–0.95). Predicted conditional occupancy probabilities of juveniles at the same 135 sites ranged from (0.35–0.92; Fig. 3c).

## Discussion

Our study assessed patterns of water loss as a process that varies spatially and temporally as a function of fine-scale environmental gradients and temporal climatic conditions. We found that spatial estimates of SAT derived from rates of water loss were significantly correlated with predicted salamander abundance and that SAT was a significant predictor of abundance as well as population demographic characteristics. Importantly, our SAT estimates were independently derived from plaster replicas deployed under field conditions, and were in no way contingent upon actual salamander distributions. Results from our study extend our understanding of plethodontid ecology and emphasize the limitations imposed by their unique physiology. Previous research has only logically conjectured the importance of hydric relations and surface activity as mechanisms underlying local distribution and population dynamics by extrapolating results from controlled laboratory experiments or indirectly through field observations (Feder 1983; Spotila 1972). As an integrated measure of the local environment and climate, SAT was a significant predictor of abundance as well as population dynamics. Combined with our findings that SAT and abundance are spatially correlated, we have compelling evidence that water loss is a physiologically-limiting factor underlying the abundance-habitat and population dynamic relationships described by Peterman and Semlitsch (2013).

Water balance can be particularly critical for smaller organisms, and we found that juvenile-sized plaster replicas lost water at 1.5–3 times greater rate than adult-sized replicas. Such differences significantly curtail surface activity, and could lead to differential survival across the landscape. In support of this, we found that the mean body size of salamanders was smaller in plots with lower rates of water loss and high SAT (Fig. 3b). Further, we found that the probability of encountering a juvenile salamander in areas of high SAT was significantly greater than areas of low SAT. In contrast, we found that adults were more uniformly distributed across the landscape, regardless of SAT (Fig. 3c). These patterns suggest that reproductive rates may be greater in high SAT regions of the landscape, or that survival of juvenile salamanders is higher in high SAT areas. Either or both of these processes would contribute to the increased abundance of salamanders in high SAT regions (Fig. 3a). Differentiating these processes as the mechanisms underlying the spatial variation in size distribution will likely only be possible through long-term, detailed studies of local demographic processes.

In corroboration with seasonal patterns of surface activity of salamanders in the field (Milanovich et al. 2006), estimated SAT differed significantly among spring and summer seasons (Table 2; Fig. S1). Although SAT was three times greater in the spring, there is still pronounced spatial heterogeneity in SAT due to the influence of topographic position in affecting water loss. The mixed effects models describing the spatial patterns of water loss for adult- and juvenile-sized replicas in the spring were nearly identical (Table 1). In the summer, juvenile replicas had no relationship with canopy cover, while adult replicas lost significantly less water as canopy cover increased. We speculate that the rate of water loss was so rapid in the high surface area juvenile models that canopy cover did little to attenuate losses. Although Peterman et al. (2013) found water loss rates of plaster replicas to be linear over an 8-hr laboratory test with up to 35% water loss, we note the possibility that rates of water loss could become non-linear as dehydration deficits approaches 100% (summer dehydration deficit for juvenile replicas: mean=60.3%, max=98.5%; adult replicas: mean= 39.4%, max=82.1%). Such non-linearity could contribute to the observed differences in parameter estimates for adult and juvenile models.

If reproductive success differs across the landscape, then *P. albagula* may best be described as existing as a spatially-structured population (Harrison 1991; Thomas and Kunin 1999). Specifically, reproductive rates and success may be greatest within forested ravines with high SAT, and be negligible or non-existent where SAT is low. As such, the presence of salamanders in low SAT areas of the landscape would predominantly depend upon salamanders dispersing from high SAT regions, implying fine-scale source-sink dynamics (Pulliam 1988). Little is known concerning dispersal in plethodontid salamanders, but as adults they are generally considered to be highly philopatric with small home ranges (Kleeberger and Werner 1982; Ousterhout and Liebgold 2010). *Plethodon cinereus* (Green), a smaller species of woodland salamander, have been found to have significant genetic differentiation over small spatial scales within continuously forested habitat (200 m; Cabe et al. 2007) and to have male-biased dispersal (Liebgold et al. 2011). Marsh et al. (2004) also found the majority of dispersing *P. cinereus* to be young adults. From a water loss perspective, smaller individuals with higher surface areas will incur the greatest cost of dispersing, so the finding of Marsh et al. (2004) that young adults are the dispersing size class may indicate a trade-off between maximizing the benefits of dispersing (e.g., reduction of kin competition, metapopulation processes, inbreeding avoidance; Hamilton and May 1977; Olivieri et al. 1995; Waser et al. 1986) while minimizing costs by not dispersing as very small, desiccation-prone juveniles. Explicit testing of how spatial variation in activity time affects population genetic structure may provide greater insight into how physiology and behaviour shape population processes.

Our findings suggest that water relations temporally and seasonally shape activity times, locally dictate habitat use, and regionally delineate distributions. Nonetheless, water loss is not a physiological process working in isolation. Metabolic rates of ectotherms are temperature dependent, increasing with environmental temperature. Because evaporative water loss also increases with temperature (Spotila 1972; Tracy et al. 2010), plethodontid salamanders are doubly challenged under hot, dry conditions. As metabolic demands increase with temperature there is a greater need for energy intake, but surface activity will likely be curtailed at higher temperatures due to increased rates of water loss. The relationship of energy expenditure and intake, as a function of temperature and foraging time (limited by water loss), was incorporated into a mechanistic energy budget model and used to accurately predict the elevational distribution of a montane woodland salamander (Gifford and Kozak 2012). Although temperature variation exists across our landscape and correlates with predicted abundance (Peterman and Semlitsch 2013), the independent (or interactive) role that spatial variation in temperature has on salamander metabolic rate, and subsequently on abundance and population dynamics, is unclear. Mechanistic modelling approaches, as used by Gifford and Kozak (2012), may be able to provide insight into these questions.

Although we observed significant spatial correlation between SAT and predicted salamander abundance, correlations were not perfect. Included in the original abundance model of Peterman and Semlitsch (2013) were topographic wetness and an interaction between topographic wetness and solar exposure. These terms were not included in our mixed effects models to limit model complexity and because there was minimal correlation with measured rates of water loss. Exclusion of these factors could explain some of the SAT-abundance discrepancies, although our mixed effects models were able to explain the majority of the variation in our data, leaving little unexplained variance to be accounted for by other factors.

Plaster replicas effectively mimicked water loss rates of living salamanders (Peterman et al. 2013), but we nonetheless made several simplifying assumptions. First, evaporative water loss in wet-skinned amphibians is determined by the moisture content of the air and the difference in the water vapour density at the surface of the animal (Spotila et al. 1992), but atmospheric moisture can vary over small spatial scales and as a function of topography and vegetation (Campbell and Norman 1998). While we attempted to account for humidity variation by using synoptic meteorological measurements, relative humidity did not correlate with water loss and was omitted from our mixed effect models. Fine-scale estimation of variation of relative humidity is likely necessary to more accurately estimate evaporative water loss in salamanders, but we note that TPI and distance from stream in our study likely correlate strongly with fine-scale humidity variation (Holden and Jolly 2011). Second, under wind-free conditions, a boundary layer will form around a stationary object (Tracy 1976), which reduces the rate of evaporative water loss. Our estimates of water loss from plaster models are therefore likely conservative as foraging or dispersal movements of surface active salamanders would disrupt the boundary layer and increase rates of water loss. Third, a critical aspect of terrestrial salamander water balance is their ability to rehydrate by absorbing water across their skin (Spotila 1972), but we sought to avoid contact of our replicas with the leaf litter and soil to minimize the potentially confounding effects of these factors on evaporative water loss.

Our study is the first to estimate spatially-explicit rates of water loss for a terrestrial amphibian under relevant ecological field conditions. Previous research has carefully detailed the physiological relationships of amphibians with their environment (reviewed by Feder 1983; Shoemaker et al. 1992; Spotila et al. 1992; Wells 2007), but only superficial attempts have been made to relate physiology with patterns observed in nature (Spotila 1972). While water loss is unlikely to be the only factor limiting terrestrial salamander activity and spatial distributions, our results provide strong support that it is critical. Future work in this system should explore how temperature, metabolic rate, and spatial energy budgets (Gifford and Kozak 2012) relate to patterns of abundance and population processes. Additionally, spatial genetic processes of terrestrial salamanders are largely unknown, but understanding how fine-scale environmental gradients relate to population and landscape genetics may provide critical insight into how physiology affects local population dynamics and dispersal. Lastly, our findings that abundance and spatial demographic patterns can be predicted by SAT have implications for the future persistence of terrestrial salamanders. Across seasons, we found that maximum temperature and time since rain were critical predictors of water loss. Current climate change scenarios are forecasting more extreme temperatures and increased variability in the interval and amount of rainfall (Field et al. 2012), and changes in these climatological parameters may profoundly affect terrestrial salamanders (Milanovich et al. 2010; Walls 2009).

By incorporating water loss and surface activity time into biophysical or dynamic population models, it may be possible to gain a better understanding of the effects that changing environmental and climatological conditions have on plethodontid salamanders.

## Acknowledgements

We thank D. Hocking for discussions on statistical analysis, and G. Connette for helpful discussion and comments. Support was provided by the University of Missouri Research Board (CB000402), Trans World Airline Scholarship, and the Department of Defense Strategic Environmental Research and Development Program (RC2155). This research was done in accordance with the laws of the state of Missouri and the USA, approved the University of Missouri Animal Care and Use Committee (#7403), and conducted under Missouri Wildlife Collector’s Permit #15203.

## Supporting Information

The following Supporting Information is available for this article online.

Appendix S1. Detailed methods of how mixed effect models were made to estimate rates of water loss.

Appendix S2. Details of field methods used to collect salamander abundance and size data, as well as detailed description of abundance and multistate modelling procedures.

Table S1. Table of Pearson’s *r* correlation coefficients for surface activity times estimated from adult- and juvenile-sized models in spring and summer.

